# Self-Supervised Pretraining for Transferable Quantitative Phase Image Cell Segmentation

**DOI:** 10.1101/2021.05.24.445397

**Authors:** Tomas Vicar, Jiri Chmelik, Roman Jakubicek, Larisa Chmelikova, Jaromir Gumulec, Jan Balvan, Ivo Provaznik, Radim Kolar

**Affiliations:** Department of Biomedical Engineering, Faculty of Electrical Engineering and Communication, Brno University of Technology, Brno, Czech Republic; Department of Pathological Physiology, Masaryk University, Brno, Czech Republic

**Keywords:** quantitative phase imaging, cell segmentation, U-Net, deep learning, self-supervised pretraining, transfer learning

## Abstract

In this paper, U-Net-based method for robust adherent cell segmentation for quantitative phase microscopy image is designed and optimised. We designed and evaluated four specific post-processing pipelines. To increase the transferability to different cell types, non-deep learning transfer with adjustable parameters is used in the post-processing step. Additionally, we proposed a self-supervised pretraining technique using nonlabelled data, which is trained to reconstruct multiple image distortions and improved the segmentation performance by from 0.67 to 0.70 of Object-wise Intersection over Union. Moreover, we publish a new dataset of manually labelled images suitable for this task together with the unlabelled data for self-supervised pretraining.

**Graphical Abstract:** 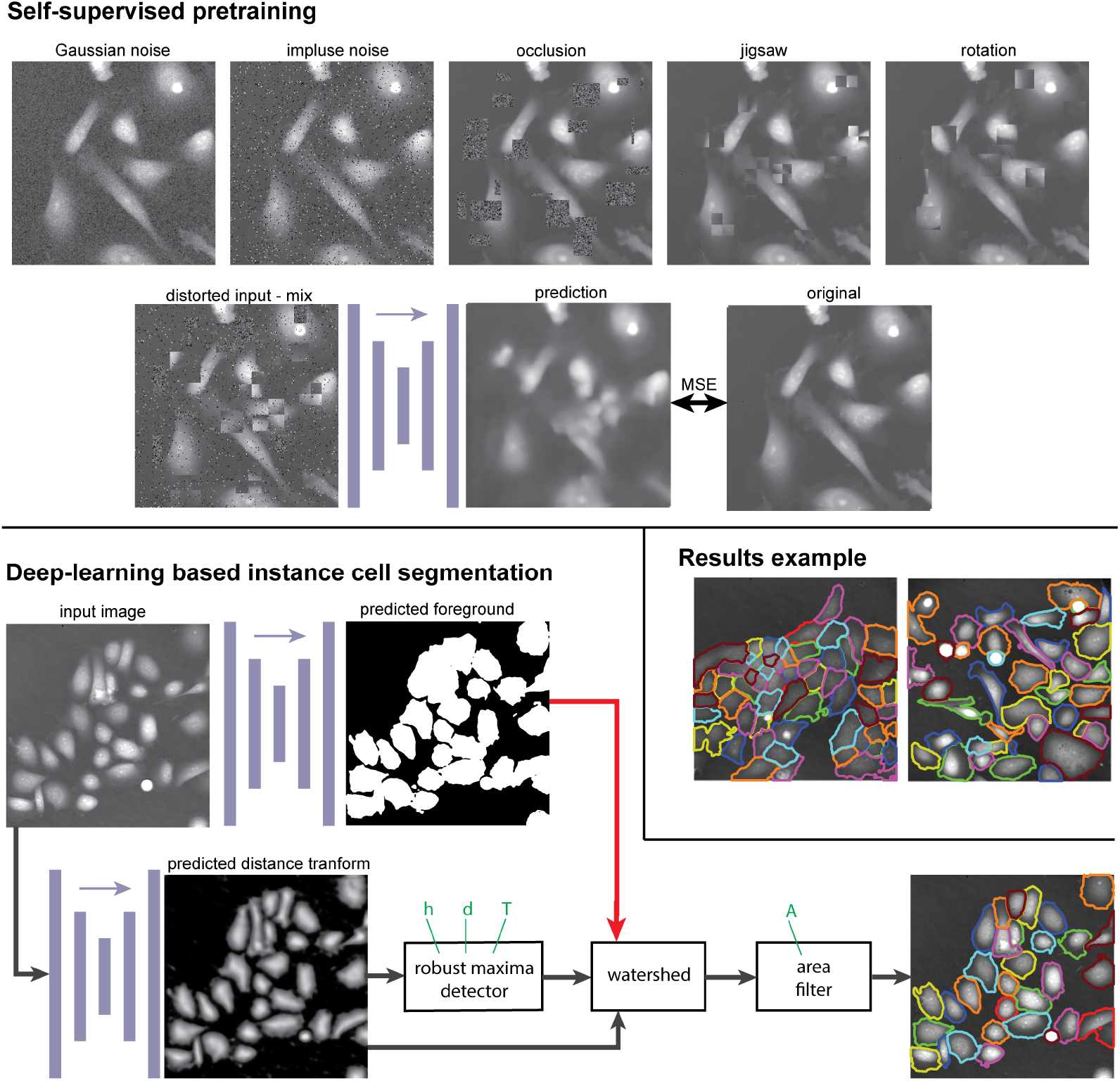

**Highlights:**

- Four strategies for instance cell segmentation with U-Net were compared.
- Specialised post-processing pipelines with tunable/optimizable parameters were designed for each segmentation strategy.
- Transferability to different cell types by optimisation of post-processing parameters was tested.
- The proposed self-supervised pretraining method improved both segmentation performance and transferability to different cell types.
- A new manually labelled quantitative phase imaging dataset for cell segmentation with unlabelled data for self-supervised pretraining was created.

## 1. Introduction

Quantitative phase imaging (QPI) has proved to be a powerful tool for label-free live cell microscopy. This technique typically provides images with superior image properties with respect to automated image processing [1]. Various QPI techniques have been developed and tested during the last decades, utilising different setups, e.g., off-axis, in-line or phase-shifting [2]. Ongoing progress in QPI microscopy enables the time-lapse observation of subtle changes in the quantitative phase dynamics of cells, such as cell dry mass distribution. It has been shown that QPI-measured dynamical changes of various parameters are typical for specific cell behaviour and can be used in different applications, e.g., cell motility assessment, homogeneity of cell content or cell mass distribution evaluation. These phase-related changes can be observed without fixation, labelling, or cell harvesting, which might severely change cell characteristics [3].

Recent papers show that instance segmentation is still critical for microscopy image segmentation in general, and QPI needs its specific setting. As we have shown [1], the cell instance segmentation based on QPI image data typically provides better results in comparison to other microscopic imaging techniques (e.g., phase contrast, differential interference contrast etc.), and relatively basic image processing techniques can provide sufficient results. However, there are still applications where precise cell segmentation is still challenging. These include populated areas of touching cells, tracking in time-lapse, and cells with complex shapes. Basic image processing methods can perform well in many cases, as shown on a combination of thresholding, hole filling, and watershed methods for yeast cell segmentation [4], Otsu-based thresholding of murine melanoma cells [5], thresholding and watershed algorithms for adherent/suspended cell classification [6], iterative thresholding method [7], or improved iterative thresholding using Laplacian of Gaussian image enhancement and distance transform-based splitting for dense cell clusters [8].

Fully convolutional neural networks (CNN), e.g., U-Net [9], with specific modifications for individual cell separation, can be successfully applied in these applications. However, direct application of U-Net for binary segmentation (foreground-background) does not achieve robust separation of individual cells because each error in boundary pixels results in the connection of these cells into one segmented cell. This can be overcome by a suitable modification of the network output, as demonstrated on various microscopic non-QPI image data [10, 11, 12, 13]. One possibility is to introduce three classes, where a ‘thicker cell boundary’ class is introduced, and, after prediction, this boundary is used to divide the cells into individual objects [10, 11]. Another simple solution is a prediction of a distance transform of the cell segmentation mask, where the foreground can be found by thresholding, and individual cells in the prediction can be found with the maxima detector. Another approach predicts the distance to a neighbouring cell or combines these multiple approaches together [12]. There is an even more complex solution of prediction of star convex polygons, where for each pixel, distance to the boundary in several directions is predicted – StarDist [13], or prediction of a vector field that can be used for cell separation – CellPose [14].

Furthermore, specific deep learning approaches have also been proposed for complex cell analysis of QPI data. Mask region-based convolution neural (Mask R-CNN [15]) network was used in two recent papers [16] [17]. A U-Net architecture [9] was also applied to QPI images, for instance, Yi et al. [18] applied U-Net to red blood cell segmentation directly on hologram images to avoid the image reconstruction part.

Recently, self-supervised pretraining methods became a popular and successful way to improve the performance of deep learning (DL) methods [19]. However, currently the best performing methods based on contrastive learning (simCLR [20] and simCLRv2 [19]) are suitable for classification tasks and not for segmentation tasks. Several other approaches for self-supervised pretraining have shown promising results, including prediction of image rotation [21], solving a jigsaw puzzle [22], image in-painting prediction [23] or denoising [24], where only the last two are suitable for segmentation networks. SeSe-Net [25] propose a more complex self-supervised approach, where two networks are trained; one is trained for the segmentation quality prediction and another for the segmentation. These two networks can then be applied for training on unlabelled data.

In this paper, we have implemented and compared four U-Net [9] based approaches for instance cell segmentation with four designed specific postprocessing pipelines. To enable the transferability of the segmentation network to different sample types without the need of annotated training data, we aimed to design these post-processing pipelines with only a few tunable parameters, which enables to perform a *non-deep learning transfer (non-DL transfer)*. Compared to standard transfer learning, this approach does not require training data and computational demanding training of DL model. We also aimed at the application of specific pretraining strategies using nonlabelled images, which can be used for self-supervised pretraining to improve final segmentation quality. The proposed methodology with self-supervised pretraining improved both segmentation performance and transferability to different cell types. Moreover, we propose a new dataset suitable for this task. Besides manually labelled data, this dataset contains unlabelled data, which can be used for self-supervised pretraining.

## 2. Material and Methods

### 2.1. Dataset

A set of adherent cell lines of various origins, tumorigenic potential, and morphology were used in this paper (PC-3, PNT1A, 22Rv1, DU145, LNCaP, A2058, A2780, A8780, Fadu, G361, HOB). PC-3, PNT1A, 22Rv1, DU145, LNCaP, A2780, and G361 cell lines were cultured in RPMI-1640 medium, A2058, FaDu, and HOB cell lines were cultured in DMEM-F12 medium, all supplemented with antibiotics (penicillin 100 U/ml and streptomycin 0.1 mg/ml), and with 10% fetal bovine serum (FBS). Prior to microscopy acquisition, the cells were maintained at 37 °C in a humidified (60%) incubator with 5% CO_2_ (Sanyo, Japan). For acquisition purposes, the cells were cultivated in the Flow chamber μ-Slide I Luer Family (Ibidi, Martinsried, Germany). To maintain standard cultivation conditions during time-lapse experiments, cells were placed in the gas chamber H201 – for Mad City Labs Z100/Z500 piezo Z-stage (Okolab, Ottaviano NA, Italy). For the acquisition of QPI, a coherence-controlled holographic microscope (Telight, Q-Phase) was used. Objective Nikon Plan 10×/0.3 was used for hologram acquisition with a CCD camera (XIMEA MR4021MC). Holographic data were numerically reconstructed with the Fourier transform method (described in [26]) and phase unwrapping was used on the phase image. QPI datasets used in this paper were acquired during various experimental setups and treatments. In most cases, experiments were conducted with the time-lapse acquisition. The final dataset contains images acquired at least three hours apart.

Our data consist of 244 labelled images of PC-3 (7,907 cells), 205 labelled PNT1A (9,288 cells) denoted as *QPI_Seg_PNT1A_PC3*, and 1,819 unlabelled images with a mixture of 22Rv1, A2058, A2780, A8780, DU145, Fadu, G361, HOB and LNCaP used for pretraining denoted as *QPI_Cell_unlabelled*. Data were labelled using a custom MATLAB semiautomatic tool, where the image is pre-segmented using, [7], and then manually edited with a set of drawing tools (e.g., cell splitting with scribble, union of selected cells, draw a new cell, delete the selected cell and correction of cell borders). An example of data is shown in Figure 4. Dataset is available at the Zenodo repository, https://doi.org/10.5281/zenodo.4771831 and the source code for semiautomatic segmentation is available together with all proposed algorithms at https://github.com/tomasvicar/Deep-QPI-Cell-Segmentation.

Labelled data were divided into training, validation, and testing sets in proportion 85/5/10% and pretraining data were divided into training and validation sets in portion 95/5%. Labelled data (training part) were also used for pretraining to make the pretraining set even larger.

### 2.2. Segmentation approaches

In this work, a novel approach for instance segmentation, inspired by [12] was designed and tested. Specifically, besides binary foreground segmentation, four other parametric images were predicted and used for splitting the foreground into individual cells. Specific post-processing (cell detection) pipelines to achieve instance cell segmentation were designed for each of these prediction approaches. A general processing scheme is shown on Figure 1a.

**Figure 1:**
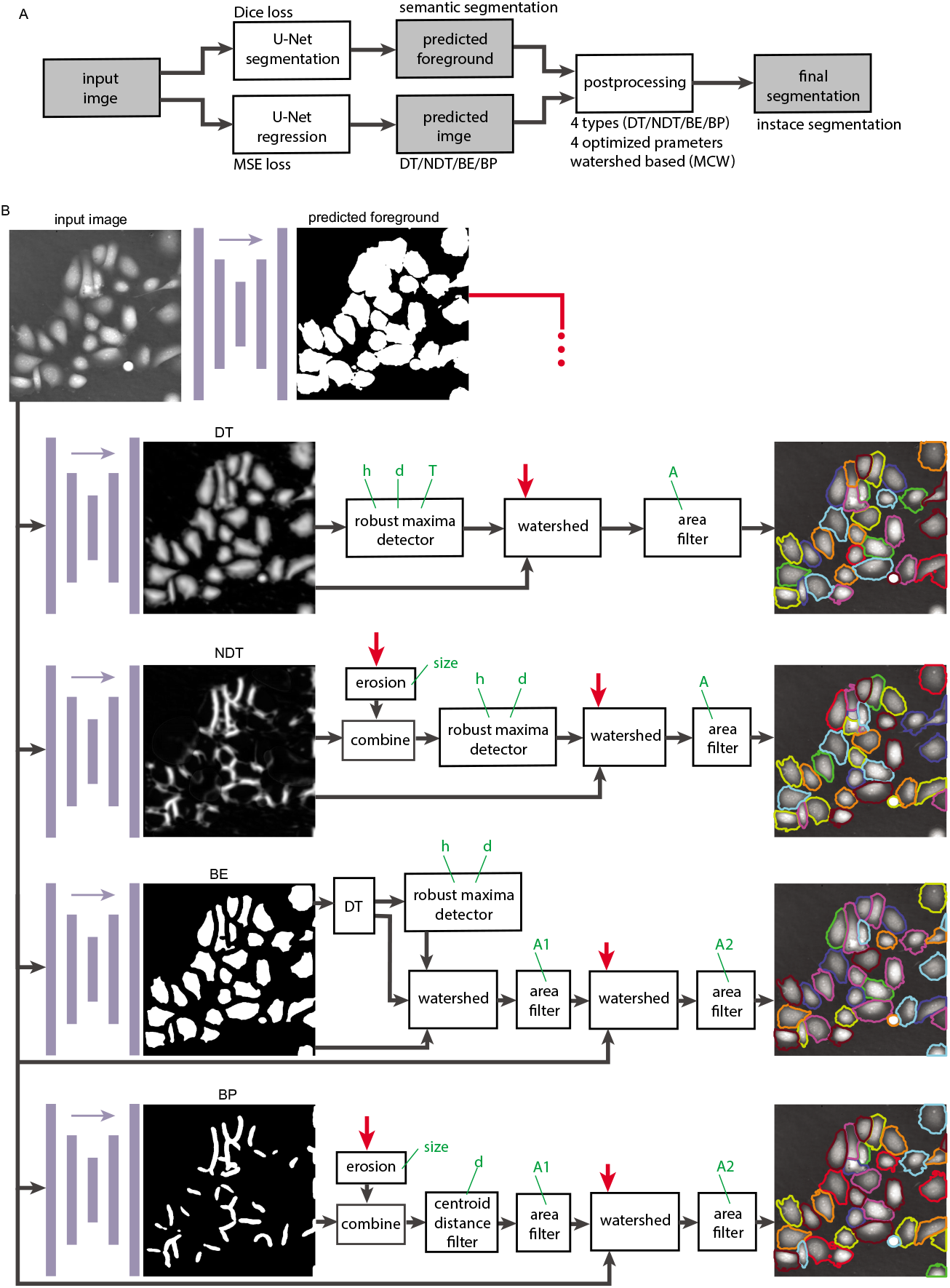
Block diagrams of tested instance segmentation methods: (a) general processing schema, (b) detailed processing scheme of individual post-processing methods (distance transform – *DT*, neighbour distance transform – *NDT*, prediction of eroded image – *BE*, prediction of boundary pixels – *BP*). Optimised parameters of the individual post-processing methods are shown in green. The red arrows indicate an input of the predicted foreground.

The U-Net [9] network with EfficientNet-B2 [27] encoder (E-U-Net) was used in our approach. Two different loss functions were used. Dice loss was used for U-Net training of pixel classification outputs. Mean Squared Error (MSE) was used for the training of U-Net with pixel regression outputs. For more details about the implementation see Appendix A.

All proposed post-processing utilise marker-controlled watershed (MCW) [28] (similarly to [11]), which is a highly efficient method in cell segmentation tasks [1] using QPI data. These post-processing approaches were further extended by subsequent steps (see Figure 1b) to make it efficient and easy to optimise by adjustment of four parameters using Bayesian optimisation [29]. In our implementation, the MCW has three inputs – (1) a binary mask (foreground mask), which is split into individual cells; (2) seeds, where every seed produce one object; (3) input image, which is used to generate the watershed borders (i.e., flooded image). Borders produced by the watershed are used to split the binary foreground mask. The last step of all methods is the area filter, which removes objects smaller than an optimized threshold. In the three post-processing pipelines, a robust maxima detector is used – it is a local maxima detector applying a constraint of the minimal distance *d* on the individual detected maxima; *h*-maxima transform [30] for a constraint of the minimal peak prominence; and threshold for minimal maxima value *T*.

Parameters of the post-processing pipeline were determined by Bayesian optimisation [29] (implementation from [31]), where the value of the cost function (Object-wise Intersection over Union, OIoU – see Section 2.4) was optimised on the validation set. The ranges of optimised parameters and the optimised values are summarized in Appendix B. A brief description of the implemented and tested pipelines follows.

#### DT

In the first approach, a normalised distance transform (*DT*) [32] image is predicted. During the training phase, this image is created from the mask of the manually segmented image and used for U-Net training. Each cell distance map is normalised to have a maximum value of one. In the inference phase, *DT* image is predicted with a trained network from the input image and used for instance cell segmentation (Figure 1b). A robust maxima detector (described above) is then applied for seed generation, and these are used together with the predicted *DT* image and the predicted foreground image as an input to the above-described MCW algorithm.

#### NDT

In the second approach, the neighbour distance transform (*NDT*) [11] is applied, and this predicted parametric map instead of the normalised DT image is used. An image transformed by *NDT* contains the values of distance to the closest cell (see Figure 1b), and background pixels are set to zero. This can be obtained with multiple DT calculations – for each cell, we can calculate DT from other cells and use the region inside this cell for *NDT*. In the post-processing phase, the predicted *NDT* image is multiplied by the eroded foreground (eroded with a circular structuring element of optimised size), and a robust maxima detector is applied to obtain seeds. Finally, *NDT* image is used as an input image for MCW.

#### BE

In the third approach, besides the foreground/background, a binary eroded (*BE*) foreground is predicted, where erosion will make a larger separation between individual cells (see Figure 1b). The amount of erosion was selected by manual tuning as a disc structuring element with a 4 pixel radius. A separation of not completely separated cells is done by the post-processing with DT, robust maxima detector (without threshold *T*) and watershed algorithm. The combination of DT with the watershed algorithm is a standard approach for splitting the connected objects in their narrowest connection (i.e., it splits the shapes in the narrowest points). With a robust maxima detector, this separation is regularised with minimal centroid distance and h-maxima transform. Output is used as a seed for the second watershed algorithm, where the negative of the original QPI image is used as the input image for MCW.

#### BP

In the fourth approach, the individual cell masks are converted into cell boundary pixels (*BP*), which can be obtained by dilatation of individual cells and determining their overlap (see Figure 1b). The amount of this dilatation was selected by manual tuning as a disc structuring element with an 8 pixel radius. In post-processing, the eroded foreground (with an optimised size structuring element) is divided by the predicted boundary pixels. The resulting seeds are filtered with an area filter and minimal distance between centroids. Besides these seeds, the negative of the original QPI image is used as the input image for the watershed.

### 2.3. Self-supervised pretraining

Self-supervised pretraining has proved to be an efficient method to improve the efficiency of different tasks in machine learning. We have tested four different approaches and their combination. All these approaches distort the input image in a different way, and the U-net is learned to restore these distortions using MSE loss. The principle of pretraining is shown in Figure 2a, and few examples of distorted images are shown on Figure 2c. The following distortions were applied:

**Additive noise with two different distributions** – impulse and Gaussian noise, where the probability of each pixel being corrupted and the standard deviation were optimised, respectively. For impulse noise, only some pixels were corrupted (with a specified probability), but with large maximal noise values of 5-times the average image standard deviation.
**Occlusion with rectangular blocks** – we have generated a set of random size rectangular blocks at random positions and replaced them with a Gaussian noise with a standard deviation equal to the average standard deviation. The number of blocks and the maximal block length of the rectangle edge were optimised. This, also known as image inpainting, is another straightforward method for pretraining of segmentation networks [23].
**Rotation of square block** – Another pretraining method of classification network is the prediction of rotation [21], which is also adapted in our application. Specifically, for rotation, we have rotated random square blocks, where the number of blocks and block size were optimised.
**Reordering of four quadrants** – Jigsaw puzzle pretraining [22] was also adapted for the segmentation network. It was implemented as a selection of random square blocks, which were split into four quadrants, and these quadrants were reshuffled. Similarly to the rotation and occlusion, the number of blocks and block size were optimised.

**Figure 2:**
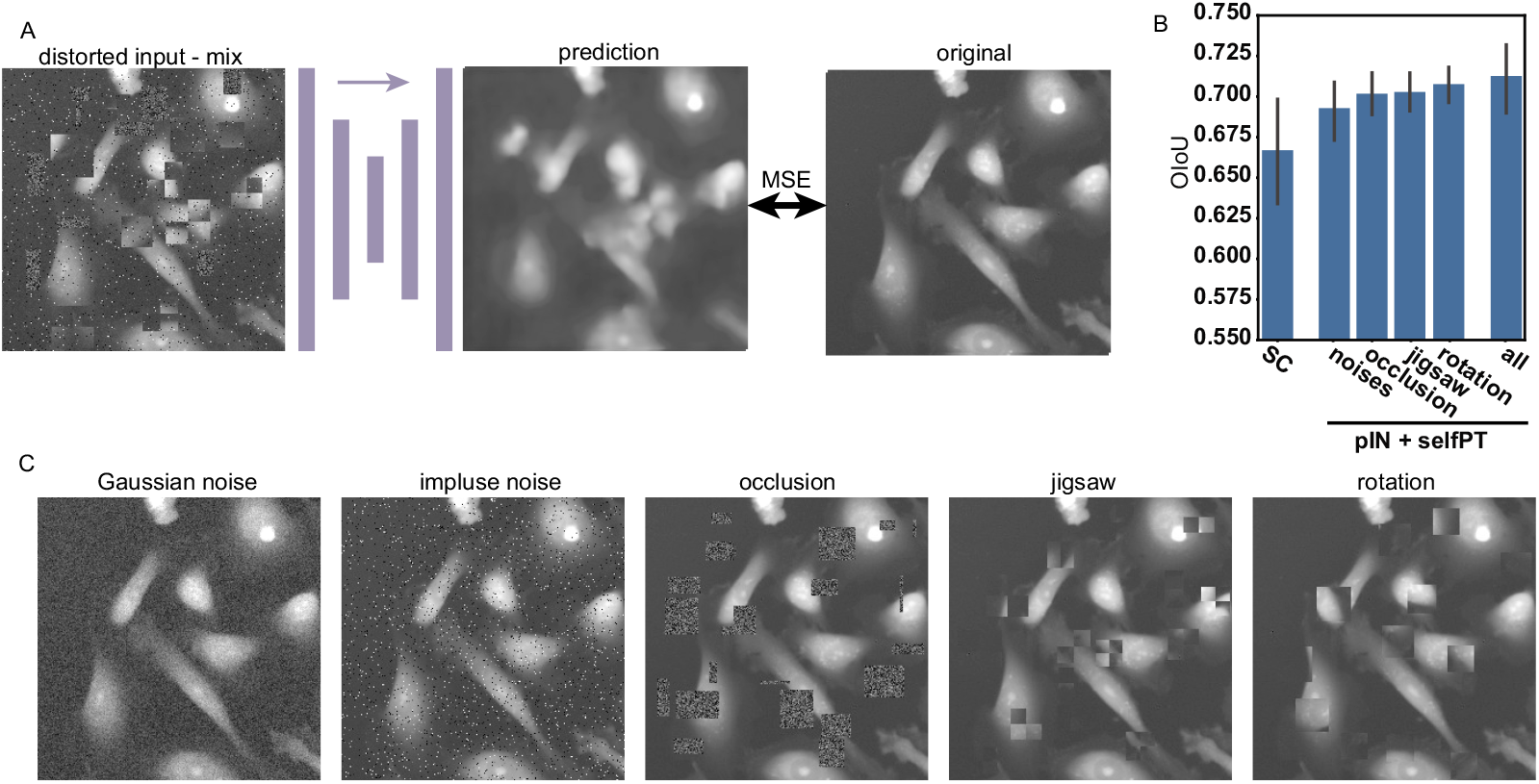
Principle and results of pretraining methods: (a) schematic example of principle of pretraining with example of network input and output, (b) Results of comparison of proposed mixed pretraining with individual image distortions used for pretraining; Distance transform (*DT*) method is used for all evaluations; for pretraining methods the network pretrained also on ImageNet beforehand (*pIN+selfPT*) is used, (c) example images of individual distortion methods Results are for 5-fold validation and bar plots show average and standard deviation.

For the final self-supervised pretraining, a mixture of all these distortions was used, where the parameters of individual distortions were optimised using Bayesian optimisation. [29], but only a single validation fold was used during optimisation. For the optimised parameters ranges and optimal values see Appendix B.

### 2.4 Evaluation metrics

For all results, the values of 5-fold validation are presented, where for each fold a new random train/validation/test split was applied. For the evaluation of semantic segmentation, the results can be easily evaluated with binary Intersection over Union (IoU):

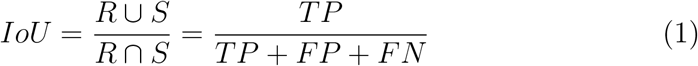

where *R* is the set of cell pixels of Ground Truth (GT) mask, *S* is the set of cell pixels in the algorithm result of semantic segmentation. *TP, FP*, and *FN* are the number of true positive, false positive, and false negative pixels, respectively. Similarly, *F*_1_ score (Dice coefficient) can be used, where it can be converted to IoU with monotonic transformation (maintaining the algorithm ranking). IoU for binary (semantic) segmentation will be denoted as BIoU.

For the evaluation of instance segmentation, an object-based metric is required. The object score *F*_1_ is defined in [10], such that the segmented cell is considered as a true positive if its IoU with the corresponding cell in the GT mask is higher than the selected threshold. A similar metric called Average Precision (AP) applies object-wise *IoU* instead of *F*_1_ score: *AP*_T_ = *TP/(TP + FP + FN)*, where subscript *T* denotes the threshold for the object to be considered as *TP* [14]. Thus, *AP* can be calculated for various thresholds with a minimum threshold of 0.5 to ensure the uniqueness of the assignment of GT cells to the resulting cell.

However, AP does not produce a single number, which would be easier to handle and would be more suitable for optimisation tasks. On the other hand, SEG score (used in the cell tracking challenge [33]) combines pixel segmentation accuracy with the correctness of identification of individual cells in a single number. In SEG, for every GT object, the segmented object with the largest IoU is found. If IoU for any GT object is smaller than 0.5, then IoU for this object is set to zero. Next, the average IoU of all GT objects is calculated. Again, the threshold 0.5 ensures that each GT object can be paired with only one segmented object. As SEG score does not contain false positive (FP) objects, we are using a more strict modification, where FP objects are counted as additional zero values for the calculation of this metric, and it will be denoted as Object-wise IoU (OIoU).

## 3. Experimental results

Several experimental setups were conducted in order to compare and test the proposed approach. First of all, U-Net based approaches were compared to non-DL approaches. Then, for the best method (post-processing pipeline), we showed the improvement achieved by the self-supervised pretraining. Afterwards, we evaluated the *non-DL transferability* of the whole pipeline to different cell types by re-optimisation of post-processing parameters only without retraining the network – non-DL-transfer.

### 3.1. Deep-learning and classical methods comparison

Comparisons of the proposed U-Net based approaches and non-DL approaches are shown on Figure 3a-c. For non-DL approaches, implementations from [3] and [8] were used. Specifically, simple threshold segmentation combined with the fast radial symmetry transform detection, *sST + dFRST*, and simple threshold segmentation combined with distance transform-based detection (*sST + dDT*) from [3] was used; improved iterative thresholding (*IIT*) from [8] and implementation of iterative thresholding by *Loewke* [7] was used. Similarly, the parameters of non-DL methods were optimised on the validation set.

**Figure 3:**
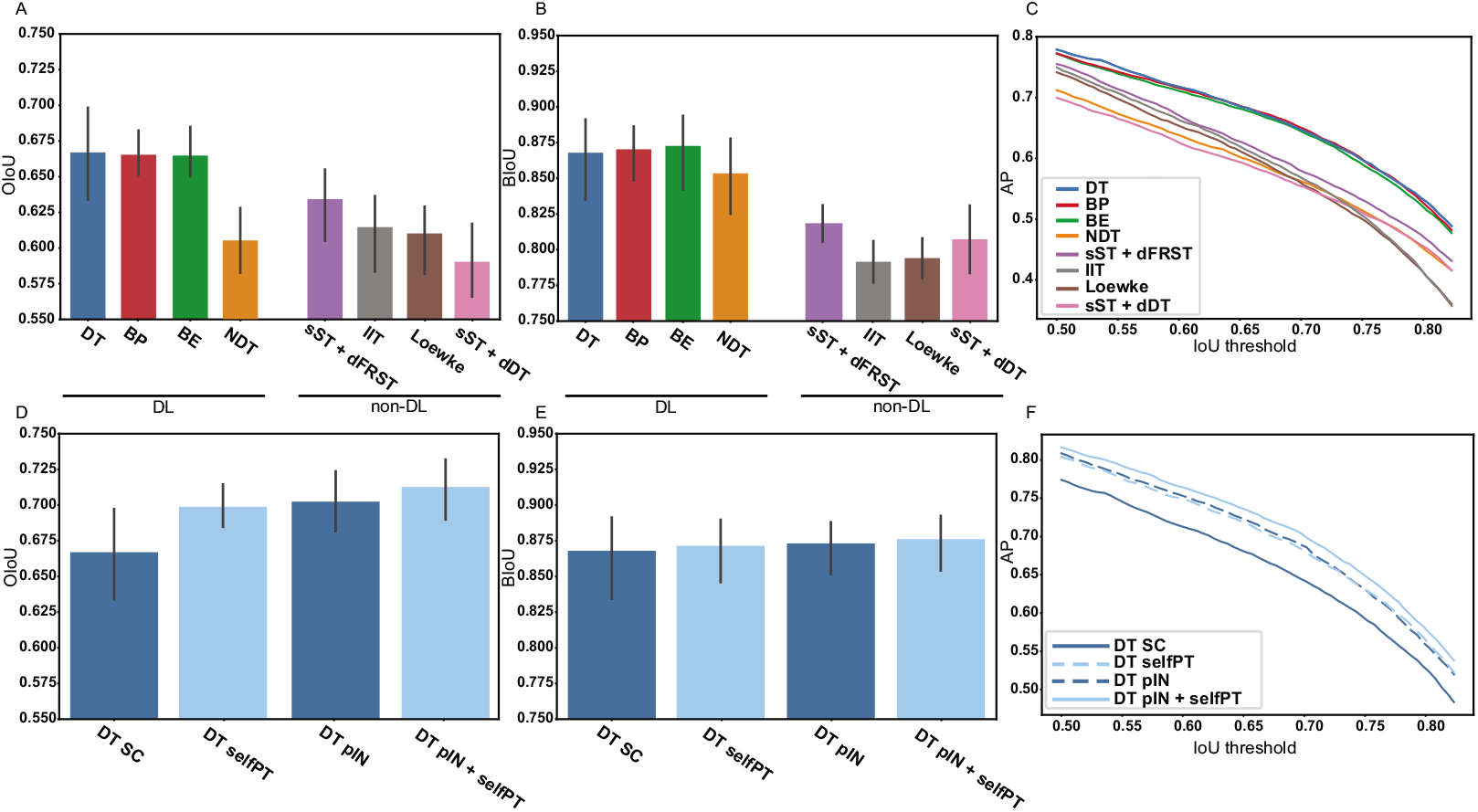
Results of the proposed methods and available non-DL method for QPI cell segmentation. Comparison of three proposed DL approaches with non-DL methods is shown in (a), (b) and (c). Comparison of proposed network trained from scratch (SC), proposed pretraining (*selfPT*), ImageNet pretrained network (*pIN*), and ImageNet pretrained network with additional proposed pretraining (*pIN+selfPT*) is shown in (d), (e) and (f), where results are shown for distance transform (*DT*) method. Results are for 5-fold validation, where AP shows average value and bar plots show average and standard deviation.

U-Net based approaches except for *NDT* method performed very similarly in all metrics (OIoU, BIoU and AP) with 0.667, 0.665 and 0.664 of OIoU for *DT, BP* and *BE* methods, respectively. Proposed *NDT* approach performed significantly worse, similarly to non-DL methods with OIoU 0.606, 0.634, 0.615, 0.610, 0.590, for *NDT, sST + dFRST, IIT, Loewke* and *sST + dDT*, respectively. They are also significantly worse in the other metrics, BIoU and AP.

Examples of results for PC-3 and PNT1A cells are shown in Figure 4. For less densely clustered cells (i.e., easier to segment), all methods performed similarly with relatively good results. However, for densely clustered cells, noticeable differences between methods can be observed. The following evaluations are performed for the best performing *DT* method in OIoU metric, which was chosen as the main optimisation metric of this paper.

**Figure 4:**
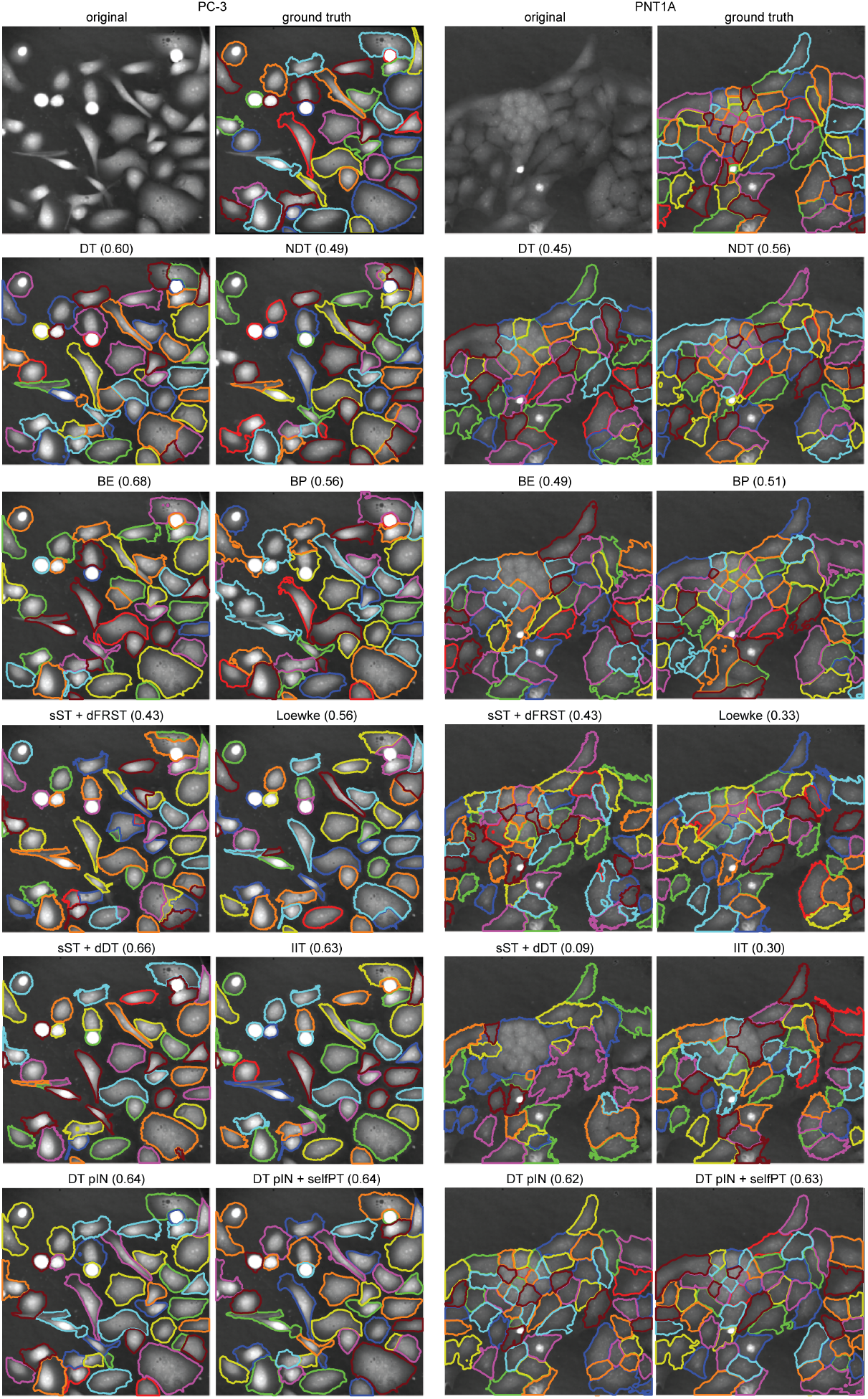
Example of results for one field of view of PNT1A and PC3 cells using different methods. The numbers represent OIoU for individual images in this example.

### 3.2. Self-supervised pretraining evaluation

Self-supervised pretraining (denoted as *selfPT*) using the optimised setting (optimised by Bayesian optimisation on the validation set) was compared to the network trained from scratch (denoted as *SC*) and the network pretrained on ImageNet dataset (denoted as *pIN*); moreover, we have tried to use ImagNet pretrained network and retrain it again using the proposed selfsupervised pretraining (denoted as *pIN+selfPT*). As shown on Figure 3d-f, *selfPT* and *pIN* networks performed similarly with OIoU 0.702 and 0.699, respectively; however, its combination (*pIN+selfPT*) leads to an additional improvement to 0.712 OIoU. The same trend is also kept for BIoU and AP. In the examples on Figure 4 can be seen that *pIN+selfPT* can significantly improve the segmentation of densely clustered cells.

Results of the comparison of the proposed mixed pretraining technique with only individual distortion pretraining are shown in Figure 2b. The best result was achieved by the proposed optimised mixture of all distortions (OIoU 0.712). The OIoU values for individual distortions were 0.693, 0.702, 0.703 and 0.708 for noise, occlusion, jigsaw, and rotation, respectively. The rotation performs best of the individual methods, and noise performs worst.

Moreover, the dependence of *SC, pIN* and *pIN + selfPT* networks on the amount of training data is shown in Figure 5, where you can see OIoU and BIoU performance of these networks for 10%, 25%, 50% and 100% of randomly selected training data. It shows that *pIN + selfPT* keeps a reasonable performance of 0.55 OIoU even in very low data regimes – 10% of training data, while *SC* and *pIN* networks failed with 0.24 OIoU and 0.31 OIoU, respectively.

**Figure 5:**
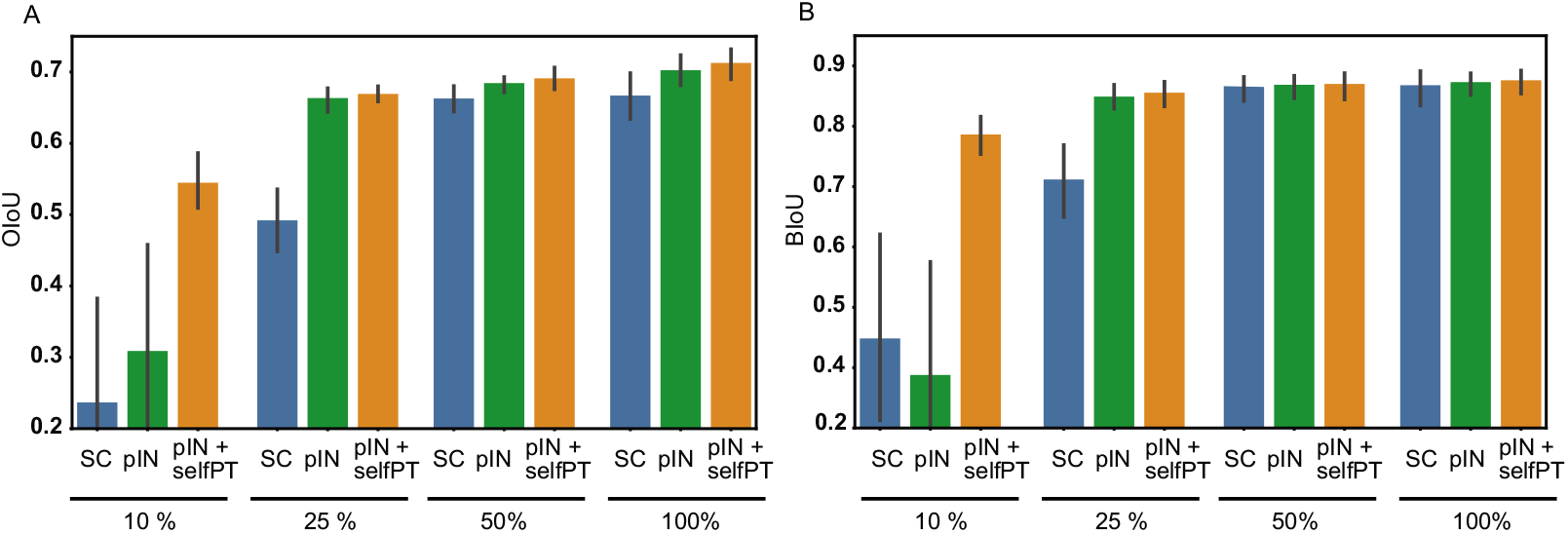
Results of the networks with reduced training set to 10%, 25%, 50% and 100% of randomly selected training data. Results of object-wise intersection over union (OIoU) and binary intersection over union (BIoU) are shown on (a) and (b), respectively. *SC*, *pIN* and *pIN + selfPT* are results for training from scratch, ImageNet pretraining and proposed self-supervised pretraining with network pretrained on ImageNet beforehand, respectively. Numbers are averages of 5-fold validation and lines in bar plots represents standard deviations.

### 3.3. Transferability with post-processing

The proposed pipeline consists of deep neural network training and postprocessing pipeline parameters optimisation, which opens the possibility of transfer to different cell lines just by adjustment of a few post-processing parameters. For this, a combination of training/optimisation/testing on PC-3/PNT1A/PC-3+PNT1A was evaluated, and the results are shown in Figure 6. Moreover, there are results of other pretraining strategies - *SC, pIN* and *pIN+selfPT*, respectively, which shows the effect of pretraining on the *non-DL transferability*.

**Figure 6:**
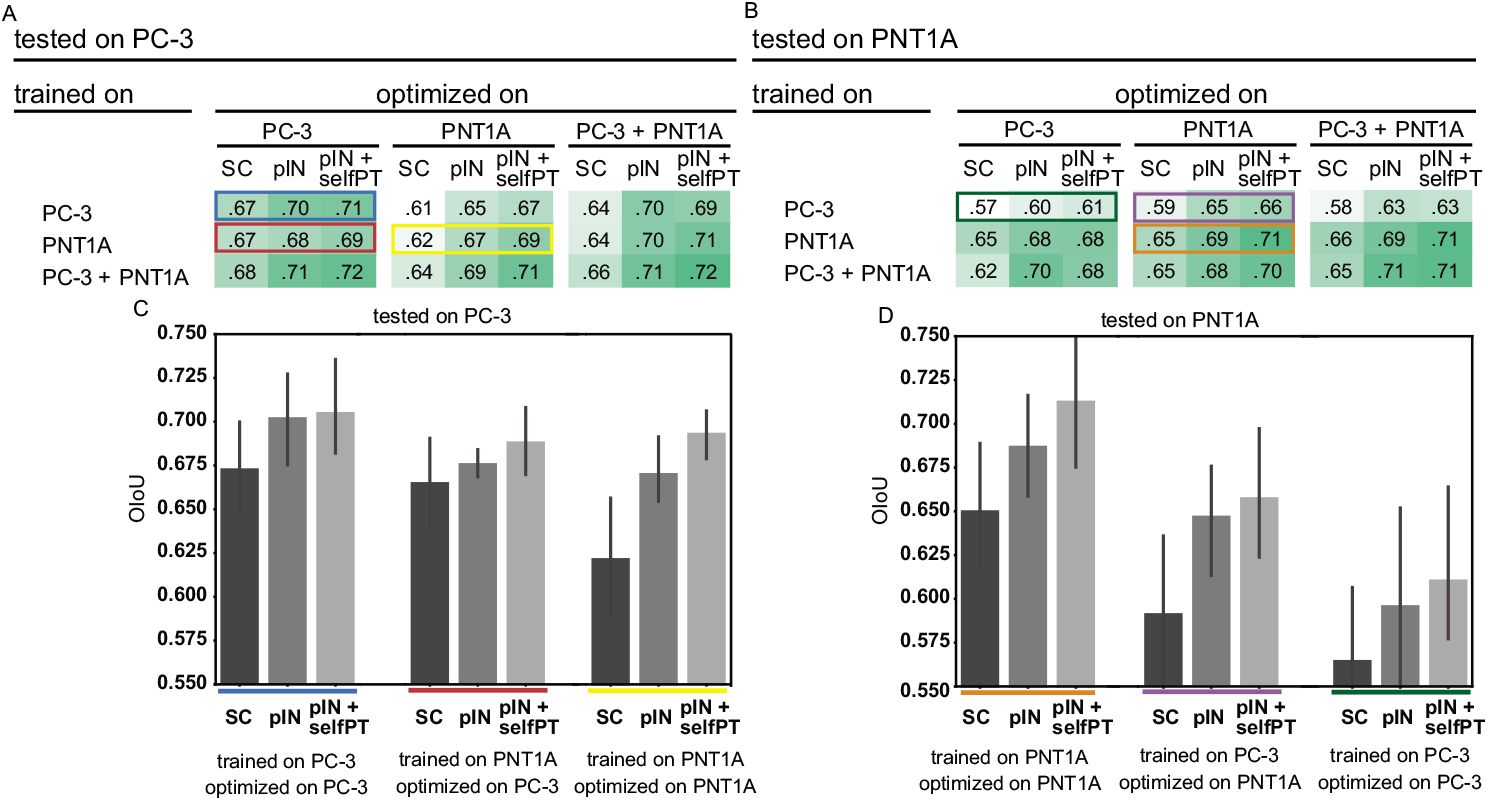
Results of transferability to different cell type by optimisation of post-processing parameters. Tables (a-b) show combinations of cell type used for training of network, cell type used for optimisation of post-processing pipeline parameters, and cell type used for evaluation. Important selected values from tables are shown on plots (c-d). *SC, pIN* and *pIN + selfPT* are results for training from scratch, ImageNet pretraining and proposed self-supervised pretraining with network pretrained on ImageNet beforehand, respectively. Numbers are averages of 5-fold validation and lines in bar plots represents standard deviations.

Segmentation results of PC-3 cells for the *pIN+selfPT* reached a very similar value for the network trained and optimised on PC-3 – 0.71 OIoU (see Figure 6c) and for the network trained on PNT1A and optimised on PC-3 – 0.69 OIoU. Even the *pIN+selfPT* network trained and optimised on PNT1A performed well on PC-3 cells – 0.69 OIoU. However, the *SC* network trained and optimised on PNT1A performed significantly worse on PC-3 cells – 0.62 OIoU.

For PNT1A cells, there are larger differences between the network trained on PNT1A and the network transferred on PNT1A by post-processing pipeline parameters optimisation. The *pIN+selfPT* network trained and optimised on PC-3 reached on PNT1A cells OIoU 0.61. When it was *non-DL transferred* by post-processing parameters optimisation (i.e., trained on PC-3 and optimised on PNT1A) it significantly improved the performance to 0.66. In comparison, the network trained and optimised using PNT1A reached 0.71. For the *SC* network trained and the *pIN* network, the same trend is kept – the OIoU performance is lower than our proposed pretraining scheme.

## 4 Discussion

The results in this paper have shown the superiority of DL methods for QPI cell segmentation over classical approaches. However, the gap between DL and non-DL approaches is not as significant as in other applications. As our results show, the network performance gradually increases with the amount of training data. The main advantage of the deep learning approach might only be evident on orders of magnitude larger datasets. However, the proposed dataset is relatively large and adding new manually segmented cells is always connected to the limited precision of a human observer. Furthermore, it must be noted that QPI is typically easily segmentable, and thus, the application of non-DL methods might provide satisfactory results, particularly for adherent cells with lower density. A benefit of DL approach arises in more difficult tasks (segmentation of high cell density with complex shapes).

In addition, a new evaluation metric named OIoU for the evaluation of instance segmentation is also proposed, which summarises the correctness of detection of individual objects together with the precision of their segmentation. Compared to AP, it produces just a single number, which can be used for the optimisation of the method; and compared to SEG score, it also penalises false positive objects.

Implementations of DL approaches in this paper were performed with two different networks – one for foreground prediction and the second for the prediction of the image with separated cells. This approach ensures avoiding ‘fight over features’ for these two tasks; however, the training of a single network for the prediction of both images together is, in principle, possible with the benefit of the faster inference.

Self-supervised pretraining (*selfPT*) on unlabelled data has shown similar performance as ImageNet pretrained network (*pIN*); however, self-supervised pretraining of ImageNet pretrained network together (*pIN+selfPT*) further increased the performance. Moreover, *pIN+selfPT* performed significantly better especially with a very small amount of training data. Different distortions applied during the pretraining have used the same framework of restoration of the distorted image, which enables to include another type of distortion. However, the combination of single distortion has led to only a negligible result improvement in comparison to patch rotation. Thus, further investigation of self-supervised pretraining may bring new findings. The self-supervised pretraining also may not be the best approach, how to efficiently utilise unlabelled data. For example, a multi-task network with self-supervised tasks and a segmentation task trained synchronously may perform even better [34]. We leave those investigations to future work.

The main advantage of optimisation in the post-processing phase is the number of optimised parameters (only four parameters in our implementation). This enables easy application of the already trained DL network to a slightly different task (i.e., different cell lines with different morphological properties) and might be a part of the solution leading to green AI strategies [35]. Moreover, these parameters can also be adjusted manually without optimisation. The success of this approach will depend on the dissimilarity of the particular tasks. However, in our case, we have shown that *non-DL transfer* is an efficient way to adjust the whole pipeline from PC-3 cell to PNT1A cell segmentation and vice versa. Furthermore, the combination with self-supervised pretraining provides an efficient way to achieve higher segmentation precision without the need for large labelled data for the new task.

## 5. Conclusion

In this paper, the U-Net-based method for robust adherent cell segmentation for quantitative phase microscopy image was designed and optimised. Four different U-Net based methods for instance cell segmentation were tested, and three of these methods achieved very similar results. These DL-based methods outperformed several well-performing non-DL methods. However, the gap between DL and non-DL methods is not so significant on a dataset of this size. Additionally, a novel self-supervised pretraining method based on image reconstruction from multiple distortions was proposed, where the proposed mixture of distortions achieved better results than each individual distortion. This improved the segmentation performance from 0.67 to 0.70 of Object-wise IoU, compared to a network trained from scratch. Another important characteristic of the proposed approach is the post-processing pipeline with adjustable parameters. This concept enables to test the non-deep learning transferability between different cell types without retraining the DL model just by optimisation of a few parameters of these post-processing pipelines. A manually segmented dataset for QPI cell segmentation (449 images) is published simultaneously with this paper (*QPI_Seg_PNT1A_PC3*) with additional unlabelled data (1,819 images) for self-supervised pretraining (*QPI_Cell_unlabelled*).

## Acknowledgment

This work is supported by the Czech Science Foundation, project no. 18-24089S. Computational resources were supplied by the project ‘e-Infrastruktura CZ’ (e-INFRA LM2018140) provided within the program Projects of Large Research, Development and Innovation Infrastructures.

## Appendix A. Implementation details

Adam optimiser was used with a decoupled weight decay [36] of 10^-5^, 1^st^ and 2^nd^ moment estimates were set to 0.9 and 0.999, respectively. Implementation of the network and ImageNet [37] pretrained weights were taken from [38]. Batch size 16 was used, where images were randomly cropped to size 256. Furthermore, augmentation with random mirroring, random affine transformation (360°rotation, max. 20% scaling, max. 5% shearing), brightness multiplying (max. 1.2×), brightness addition (max. ±0.2), blurring/sharpening (max. 0.5 Gaussian sigma and max. subtraction of 0.5 × Laplacian). Initial learning rate 0.01 decaying to 10% of its previous value each 40, 20, 10 and 5 epochs was used.

## Appendix B. Optimised parameters

**Table B.1:** Table of optimised parameters for pretraining. Range is range of values searched with Bayesian optimisation. Optimal values is for *DTpIN+selfPT* method (distance transform based method with ImageNet pretrained network and with additional self supervised pretraining). Blocks for occlusion, jigsaw and rotation were generated in random order for 256 × 256 image patch, where block was skipped, if it overlaps with previous blocks.

| distortion | parameter | range | optimal |
| --- | --- | --- | --- |
| noises | Gaussian noise sigma (fraction of image std) | 0 – 1 | 0.76 |
|  | impulse noise probability | 0 – 0.5 | 0.19 |
| occlusion | rectangle edge max length | 10 – 70 | 39 |
|  | number of rectangles | 0 – 50 | 9 |
| jigsaw | square edge max length | 10 – 70 | 40 |
|  | number of squares | 0 – 50 | 11 |
| rotation | square edge max length | 10 – 70 | 35 |
|  | number of squares | 0 – 50 | 8 |

**Table B.2:** Table of optimised parameters of individual methods. Ranges of values searched with Bayesian optimisation together with optimal values for individual folds within 5-fold validation are shown.

| method | parameters | range | fold |  |  |  |  |
| --- | --- | --- | --- | --- | --- | --- | --- |
|  |  |  | 1 | 2 | 3 | 4 | 5 |
| DT | h - h-maxima | 0.05 - 0.9 | 0.17 | 0.21 | 0.14 | 0.21 | 0.20 |
|  | d - minimal distance | 1 - 80 | 12 | 2 | 1 | 3 | 11 |
|  | T - threshold of DT | 0 - 0.9 | 0.46 | 0.58 | 0.62 | 0.42 | 0.62 |
|  | A - area filter | 20 - 400 | 95 | 91 | 83 | 82 | 103 |
| NDT | size - erosion radius | 1 - 20 | 11 | 11 | 11 | 11 | 11 |
|  | h - h-maxima | 0.05 - 0.9 | 0.11 | 0.09 | 0.13 | 0.11 | 0.12 |
|  | d - minimal distance | 1 - 80 | 8 | 8 | 8 | 9 | 8 |
|  | A - area filter | 20 - 400 | 250 | 201 | 222 | 185 | 225 |
| BE | h - h-maxima | 0.05 - 10 | 2 | 9 | 9 | 0 | 10 |
|  | d - minimal distance | 1 - 100 | 95 | 85 | 99 | 100 | 83 |
|  | A1 - area filter1 | 10 - 500 | 10 | 30 | 71 | 70 | 63 |
|  | A2 - area filter2 | 10 - 500 | 500 | 406 | 267 | 472 | 457 |
| BP | size - erosion radius | 0 - 6 | 6 | 5 | 5 | 5 | 6 |
|  | d - minimal distance | 1 - 100 | 5 | 2 | 1 | 5 | 7 |
|  | A1 - area filter1 | 10 - 500 | 10 | 42 | 78 | 36 | 81 |
|  | A2 - area filter2 | 10 - 500 | 441 | 458 | 475 | 497 | 500 |
| sST+dDT | foreground threshold | -0.2 - 0.25 | 0.03 | 0.03 | 0.05 | 0.03 | 0.03 |
|  | foreground area filter | 5 - 400 | 303 | 148 | 391 | 280 | 36 |
|  | foreground hole filter | 5 - 250 | 130 | 115 | 190 | 71 | 79 |
|  | final area filter | 5 - 400 | 237 | 257 | 210 | 62 | 43 |
|  | final hole filter | 5 - 250 | 115 | 50 | 51 | 75 | 112 |
|  | h-maxima for DT | 0 - 50 | 1.27 | 1.87 | 1.08 | 1.13 | 1.24 |
|  | threshold for DT | 0 - 1.2 | 0.31 | 0.28 | 0.25 | 0.36 | 0.28 |
| sST+dFRST | foreground threshold | -0.2 - 0.25 | 0.03 | 0.03 | 0.03 | 0.03 | 0.03 |
|  | area filter | 5 - 400 | 378 | 210 | 329 | 350 | 395 |
|  | FRST threshold | 0 - 0.7 | 0.35 | 0.20 | 0.48 | 0.63 | 0.49 |
|  | FRST k | 0.5 - 30 | 1.29 | 1.86 | 1.30 | 0.95 | 0.94 |
|  | FRST alfa | 0.01 - 10 | 7.88 | 1.33 | 5.98 | 5.89 | 6.19 |
|  | FRST min. radius | 2 - 35 | 10.73 | 10.79 | 10.02 | 8.45 | 8.49 |
|  | FRST max. radius | 8 - 90 | 42.39 | 50.80 | 58.65 | 73.57 | 27.18 |
| Loewke | minimal mass | 20 - 200 | 114 | 126 | 127 | 123 | 91 |
|  | hole filter | 10 - 100 | 99 | 65 | 99 | 99 | 19 |
| IIT | sigma | 0.02 - 20 | 0.06 | 0.02 | 0.06 | 0.02 | 0.03 |
|  | lambda | 1 - 30 | 2.16 | 9.09 | 26.40 | 5.06 | 27.81 |
|  | minimal mass | 10 - 400 | 240 | 131 | 180 | 134 | 112 |
|  | h-maxima (h) | 0 - 10 | 2.45 | 8.64 | 2.97 | 2.11 | 3.00 |

## References

[1] T. Vicar, J. Balvan, J. Jaros, F. Jug, R. Kolar, M. Masarik, J. Gumulec, Cell segmentation methods for label-free contrast microscopy: review and comprehensive comparison, BMC bioinformatics 20 (360) (2019) 1–25. doi:10.1186/s12859-019-2880-8.

[2] M. K. Kim, Principles and techniques of digital holographic microscopy, SPIE Reviews 1 (1) (2010) 1–51. doi:10.1117/6.0000006.

[3] T. Vicar, M. Raudenska, J. Gumulec, J. Balvan, The quantitative-phase dynamics of apoptosis and lytic cell death, Scientific reports 10 (1) (2020) 1–12. doi:10.1038/s41598-020-58474-w.

[4] H. Alanazi, A. J. Canul, A. Garman, J. Quimby, A. E. Vasdekis, Robust microbial cell segmentation by optical-phase thresholding with minimal processing requirements, Cytometry Part A 91 (5) (2017) 443–449. doi:doi.org/10.1002/cyto.a.23099.

[5] V. L. Calin, M. Mihailescu, N. Tarba, A. M. Sandu, E. Scarlat, M. G. Moisescu, T. Savopol, Digital holographic microscopy evaluation of dynamic cell response to electroporation, Biomed. Opt. Express 12 (4) (2021) 2519–2530. doi:10.1364/BOE.421959.

[6] B. Kemper, H. Eilers, T. Klein, K. Brinker, S. Ketelhut, Quantitative phase imaging-based machine learning approaches for the analysis of adherent and suspended cells, in: T. G. Brown, T. Wilson, L. Waller (Eds.), Three-Dimensional and Multidimensional Microscopy: Image Acquisition and Processing XXVIII, Vol. 11649, International Society for Optics and Photonics, SPIE, 2021, pp. 22–27. doi:10.1117/12.2577825.

[7] N. O. Loewke, S. Pai, C. Cordeiro, D. Black, B. L. King, C. H. Contag, B. Chen, T. M. Baer, O. Solgaard, Automated cell segmentation for quantitative phase microscopy, IEEE transactions on medical imaging 37 (4) (2017) 929–940. doi:10.1109/TMI.2017.2775604.

[8] T. Vicar, J. Chmelik, R. Kolar, Cell segmentation in quantitative phase images with improved iterative thresholding method, in: 8th European Medical and Biological Engineering Conference, Springer International Publishing, Cham, 2021, pp. 233–239. doi:10.1007/978-3-030-64610-3_27.

[9] O. Ronneberger, P. Fischer, T. Brox, U-Net: Convolutional networks for biomedical image segmentation, in: N. Navab, J. Hornegger, W. Wells, A. Frangi (Eds.), Medical Image Computing and Computer-Assisted Intervention 2015 (MICCAI 2015), Vol. 9351 of Lecture Notes in Computer Science, Springer, 2015, pp. 234–241. doi:10.1007/978-3-319-24574-4_28.

[10] J. C. Caicedo, J. Roth, A. Goodman, T. Becker, K. W. Karhohs, M. Broisin, C. Molnar, C. McQuin, S. Singh, F. J. Theis, et al., Evaluation of deep learning strategies for nucleus segmentation in fluorescence images, Cytometry Part A 95 (9) (2019) 952–965. doi:10.1002/cyto.a.23863.

[11] F. Lux, P. Matula, Cell segmentation by combining marker-controlled watershed and deep learning, arXiv preprint arXiv:2004.01607 (2020).

[12] T. Scherr, K. Löffler, M. Böhland, R. Mikut, Cell segmentation and tracking using distance transform predictions and movement estimation with graph-based matching, arXiv preprint arXiv:2004.01486 (2020).

[13] U. Schmidt, M. Weigert, C. Broaddus, G. Myers, Cell detection with star-convex polygons, in: A. F. Frangi, J. A. Schnabel, C. Davatzikos, C. Alberola-López, G. Fichtinger (Eds.), Medical Image Computing and Computer-Assisted Intervention 2018 (MICCAI 2018), Vol. 11071 of Lecture Notes in Computer Science, Springer, 2018, pp. 265–273. doi:10.1007/978-3-030-00934-2_30.

[14] C. Stringer, T. Wang, M. Michaelos, M. Pachitariu, Cellpose: a generalist algorithm for cellular segmentation, Nature Methods 18 (1) (2021) 100–106. doi:10.1038/s41592-020-01018-x.

[15] K. He, G. Gkioxari, P. Dollár, R. Girshick, Mask r-cnn, in: 2017 IEEE International Conference on Computer Vision (ICCV), IEEE, 2017, pp. 2961–2969. doi:10.1109/ICCV.2017.322.

[16] K. Eder, T. Kutscher, A. Marzi, Alvaro Barroso, J. Schnekenburger, B. Kemper, Automated detection of macrophages in quantitative phase images by deep learning using a mask region-based convolutional neural network, in: N. T. Shaked, O. Hayden (Eds.), Label-free Biomedical Imaging and Sensing (LBIS) 2021, Vol. 11655, International Society for Optics and Photonics, SPIE, 2021, pp. 88–94. doi:10.1117/12.2577232.

[17] Y.-H. Lin, K. Y.-K. Liao, K.-B. Sung, Automatic detection and characterization of quantitative phase images of thalassemic red blood cells using a mask region-based convolutional neural network, Journal of Biomedical Optics 25 (11) (2020) 1–14. doi:10.1117/1.JBO.25.11.116502.

[18] F. Yi, I. Moon, B. Javidi, Automated red blood cells extraction from holographic images using fully convolutional neural networks, Biomed. Opt. Express 8 (10) (2017) 4466–4479. doi:10.1364/BOE.8.004466.

[19] T. Chen, S. Kornblith, K. Swersky, M. Norouzi, G. Hinton, Big selfsupervised models are strong semi-supervised learners, arXiv preprint arXiv:2006.10029 (2020).

[20] T. Chen, S. Kornblith, M. Norouzi, G. Hinton, A simple framework for contrastive learning of visual representations, in: International conference on machine learning, PMLR, 2020, pp. 1597–1607.

[21] S. Gidaris, P. Singh, N. Komodakis, Unsupervised representation learning by predicting image rotations, arXiv preprint arXiv:1803.07728 (2018).

[22] C. Doersch, A. Gupta, A. A. Efros, Unsupervised visual representation learning by context prediction, in: Proceedings of the IEEE international conference on computer vision, 2015, pp. 1422–1430. doi:10.1109/ICCV.2015.167.

[23] D. Pathak, P. Krahenbuhl, J. Donahue, T. Darrell, A. A. Efros, Context encoders: Feature learning by inpainting, in: Proceedings of the IEEE conference on computer vision and pattern recognition, 2016, pp. 2536–2544. doi:10.1109/CVPR.2016.278.

[24] M. Prakash, T.-O. Buchholz, M. Lalit, P. Tomancak, F. Jug, A. Krull, Leveraging self-supervised denoising for image segmentation, in: 2020 IEEE 17th International Symposium on Biomedical Imaging (ISBI), IEEE, 2020, pp. 428–432. doi:10.1109/ISBI45749.2020.9098559.

[25] Z. Zeng, Y. Xulei, Y. Qiyun, Y. Meng, Z. Le, Sese-net: Self-supervised deep learning for segmentation, Pattern Recognition Letters 128 (2019) 23–29. doi:10.1016/j.patrec.2019.08.002.

[26] T. Slaby, P. Kolman, Z. Dostal, M. Antos, M. Lostak, R. Chmelik, Off-axis setup taking full advantage of incoherent illumination in coherence-controlled holographic microscope, Optics Express 21 (12) (2013) 14747–14762. doi:10.1364/OE.21.014747.

[27] M. Tan, Q. Le, Efficientnet: Rethinking model scaling for convolutional neural networks, in: International Conference on Machine Learning, PMLR, 2019, pp. 6105–6114.

[28] F. Meyer, Topographic distance and watershed lines, Signal processing 38 (1) (1994) 113–125. doi:10.1016/0165-1684(94)90060-4.

[29] J. Snoek, H. Larochelle, R. P. Adams, Practical bayesian optimization of machine learning algorithms, in: Advances in neural information processing systems, 2012, pp. 2951–2959. doi:10.5555/2999325.2999464.

[30] K. Thirusittampalam, M. J. Hossain, O. Ghita, P. F. Whelan, A novel framework for cellular tracking and mitosis detection in dense phase contrast microscopy images, IEEE journal of biomedical and health informatics 17 (3) (2013) 642–653. doi:10.1109/titb.2012.2228663.

[31] F. Nogueira, Bayesian Optimization: Open source constrained global optimization tool for Python (2014–). URL https://github.com/fmfn/BayesianOptimization

[32] C. R. Maurer, R. Qi, V. Raghavan, A linear time algorithm for computing exact euclidean distance transforms of binary images in arbitrary dimensions, IEEE Transactions on Pattern Analysis and Machine Intelligence 25 (2) (2003) 265–270. doi:10.1109/TPAMI.2003.1177156.

[33] V. Ulman, M. Maška, K. E. Magnusson, O. Ronneberger, C. Haubold, N. Harder, P. Matula, P. Matula, D. Svoboda, M. Radojevic, et al., An objective comparison of cell-tracking algorithms, Nature methods 14 (12) (2017) 1141–1152. doi:10.1038/nmeth.4473.

[34] S. Reiß, C. Seibold, A. Freytag, E. Rodner, R. Stiefelhagen, Every annotation counts: Multi-label deep supervision for medical image segmentation, arXiv preprint arXiv:2104.13243 (2021).

[35] R. Schwartz, J. Dodge, N. A. Smith, O. Etzioni, Green ai, Communications of the ACM 63 (2020) 54–63. doi:10.1145/3381831.

[36] I. Loshchilov, F. Hutter, Decoupled weight decay regularization, arXiv preprint arXiv:1711.05101 (2017).

[37] J. Deng, W. Dong, R. Socher, L.-J. Li, K. Li, L. Fei-Fei, Imagenet: A large-scale hierarchical image database, in: 2009 IEEE conference on computer vision and pattern recognition, IEEE, 2009, pp. 248–255. doi:10.1109/CVPR.2009.5206848.

[38] P. Yakubovskiy, Segmentation models pytorch, https://github.com/qubvel/segmentation_models.pytorch (2020).

